# Spatial epigenomics reveals ancestry-associated regulatory states linked to aggressive prostate cancer biology

**DOI:** 10.64898/2026.10.16.753149

**Authors:** Beatriz German, Kun-Lin Ho, Jennifer Garbarino, Sadia Sayed, Amina Ali, William Azadze, Tri Truong, Sally Elsamanoudi, James McGann, Raunak Shrestha, Colin Ng, Craig D. Shriver, Gregory Chesnut, Leigh Ellis

**Author notes:** Corresponding Author: Dr. Beatriz German and Dr. Leigh Ellis, and. Co-first authors (Beatriz German, Kun-Lin Ho).

## Abstract

Prostate cancer (PCa) exhibits substantial biological heterogeneity, including population associated differences in tumor aggressiveness and clinical outcomes. Although socioeconomic and environmental factors contribute to these disparities, intrinsic tumor regulatory programs may also influence disease behavior. Here, we investigated whether differences in chromatin regulation contribute to distinct tumor states in Black and White men with localized PCa. We performed spatial chromatin accessibility profiling (ATAC-seq) of prostatectomy specimens from Black and White patients receiving equal healthcare access at the Center for Prostate Disease Research (USUHS) and Walter Reed National Military Medical Center. Tumors showed distinct chromatin accessibility landscapes between the two populations, with tumors from Black patients exhibiting reduced accessibility of motifs associated with the chromatin architectural regulator CTCF and enrichment of motifs linked to epithelial-mesenchymal transition and inflammatory programs. Reduced CTCF expression was independently validated by immunohistochemistry in 99 prostatectomy specimens. These findings identify population-associated differences in chromatin regulation in localized PCa and suggest that altered CTCF-associated regulation, inflammatory signaling, and epithelial plasticity may contribute to the biological heterogeneity underlying ancestry-dependent PCa disparities.

## Introduction

Prostate cancer (PCa) represents one of the most significant examples of racial disparities in cancer biology. In the United States, Black men are approximately 1.7 times more likely to develop PCa and more than two times more likely to die from the disease compared with White men. However, studies conducted in equal access healthcare settings have yielded mixed findings. Several analyses within the Department of Defense Military Health System and Veterans Health Administration have reported comparable survival, treatment patterns, or disease progression between Black and White men (1–3), whereas others have identified persistent differences, including higher PCa incidence and metastatic disease at diagnosis and shorter time to biochemical recurrence after radical prostatectomy among Black men (4–8). Thus, equal access to healthcare may attenuate some disparities but does not uniformly eliminate population associated differences in PCa disease course.

The biological basis of these differences remains incompletely understood. Although germline variants and somatic mutations contribute to PCa susceptibility and progression (9, 10), epigenetic mechanisms, including DNA methylation, chromatin accessibility, histone modifications, and non-coding regulatory element activity, are also important determinants of tumor phenotype and disease progression (11–13). In PCa, lineage factors such as androgen receptor (AR) signaling, lineage specification, phenotypic plasticity, immune response, cell cycle control (14–17). In PCa, chromatin organization shaped by lineage factors such as AR, FOXA1 and HOXB13 regulates tumor-cell identity and plasticity. During progression, remodeling of this regulatory landscape can enable transitions toward aggressive stem-like, EMT, and neuroendocrine states, particularly following loss independence of canonical AR activity (16, 18, 19). Notably, AP-1 activity is a key feature of AR-low/stem-like PCa and contributes to alternative regulatory programs associated with tumor plasticity (20, 21).

Consistent with a role for epigenetic regulation in population associated tumor biology, multiple studies have identified differences in epigenome and transcriptional regulatory landscape of PCa across ancestry groups. DNA methylation profiling from The Cancer Genome Atlas (TCGA) revealed differential methylation landscapes between tumors from Black and White patients linked to tumor progression and recurrence (22)(23)(24). Transcriptional studies have further identified differences in inflammatory signaling, androgen receptor (AR) signaling, estrogen receptor pathways, and chromatin regulation (25–27). Importantly, it has been demonstrated that the AR cistrome differs between Black and White patients, revealing ancestry-associated differences in AR binding landscapes and regulatory programs that influence transcriptional output (27). These findings highlight enhancer organization and transcription factor activity as key regulators of population-associated differences in PCa biology. Recurrent genomic alterations such as TMPRSS2–ERG fusions and PTEN loss also occur at different frequencies across ancestry groups (28), suggesting that distinct regulatory contexts may contribute to tumor evolution. Studies of aggressive PCa in Black men have identified differences in inflammatory, metabolic, and epigenetic programs that may contribute to population associated tumor aggressiveness (29, 30). Together, these findings support a role for regulatory state in population associated PCa heterogeneity, but most studies have relied on bulk profiling and therefore provide limited resolution of how these programs are distributed across tumor cells and tissue niches.

Whether ancestry-associated differences in chromatin regulation define distinct tumor cell states remain unclear. In particular, it is not known how chromatin accessibility is organized across malignant epithelial regions or whether regulatory elements associated with epithelial identity plasticity, and alternative transcriptional states differ between ancestry groups. This question is relevant to the interplay between chromatin architecture and transcription factor activity. Resolving the chromatin regulatory features within their spatial context may therefore provide insight into how chromatin states relate to tumor cell phenotypes.

Recent advances in single-cell and spatial epigenomics now enable genome-wide profiling of chromatin accessibility while preserving cellular identity and tissue architecture (31, 32). These approaches have revealed previously unrecognized cellular states, enhancer programs, and regulatory circuits driving tumor evolution, therapeutic resistance, and phenotypic plasticity across different tissue types (33). Here, we used spatial chromatin accessibility profiling to characterize primary prostate tumors from Black and White patients treated within the Military Health System. We identified distinct chromatin accessibility landscapes, including reduced accessibility of CCCTC-binding factor (CTCF) motifs in tumors from Black patients, together with enrichment of regulatory programs associated with epithelial plasticity and inflammatory signaling. Together, these findings identify population associated differences in chromatin regulation within localized PCa and provide a framework for understanding how tumor regulatory states contribute to biological heterogeneity.

## Results

### Overview and quality metrics of Spatial ATAC-seq in Race-stratified Prostate Cancer Biospecimens

To define ancestry-associated chromatin landscape in primary prostate cancer (PCa), we performed spatial ATAC-seq on treatment-naïve prostatectomy specimens from 10 patients (five White and five Black) collected through the Center for Prostate Disease Research (CPDR), Walter-Reed Department of Surgery (**Fig. 1A**). All the samples were matched as closely as possible for age (median, 60.4 years), Gleason score (median, 7), and serum PSA levels (median, 6.15 ng/mL) to minimize potential confounding effects (clinical characteristics are summarized in **Fig. 1B**). The spatial ATAC-seq workflow is outlined in **Fig. 1C**. Briefly, tissue sections underwent Tn5 transposition followed by spatial barcoding to generate spatially resolved chromatin accessibility profiles. Consecutive tissue sections were subjected to H&E staining for histopathological annotation and spatial alignment of chromatin accessibility data (**Fig. S2A**).

**Figure 1:**
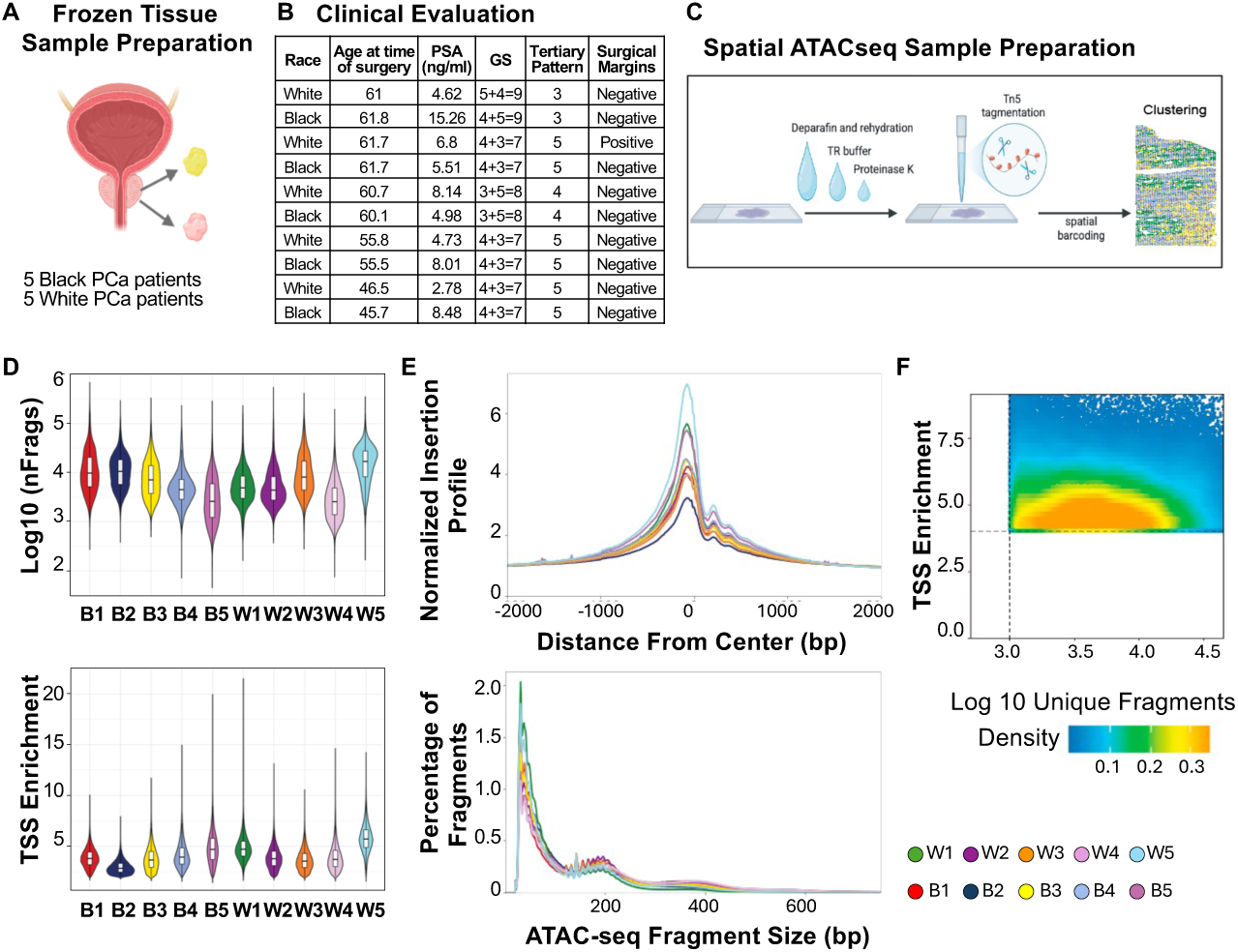
Overview and Quality Metrics of Spatial ATAC-seq in Race-Stratified Prostate Cancer Biospecimens. **A.** Schematic overview of the spatial ATAC-seq study design. Radical prostatectomy specimens were collected from patients with primary prostate cancer (PCa), including Black (n = 5) and White (n = 5) individuals. **B.** Clinical and pathological characteristics of the patient cohort, as assessed by a board-certified pathologist. **C.** Overview of the Spatial ATAC-seq workflow. Tissue sections were subjected to Tn5 transposition followed by in situ ligation with two sets of spatially resolved DNA barcodes to generate spatially indexed chromatin accessibility profiles. **D.** Quality control metrics showing the number of unique fragments per sample (top) and the fraction of reads mapping to transcription start sites (TSS) (bottom). **E**. Insert size distribution of ATAC-seq fragments across samples (top) and enrichment of reads around TSSs (bottom). **F.** Relationship between TSS enrichment score and the number of unique nuclear fragments per cell across all samples.

Across the dataset, spatial ATAC-seq yielded a media of 10,953 Fragments per 12μm spot (**Fig. S2B-C**), sufficient chromatin accessibility coverage across the dataset. The transcription starts site (TSS) enrichment scores averaged 4.0 (**Fig. 1D**) and the fraction of reads in peaks (FRIP) exceeded 0.2 across all samples (**Fig. S2B-C**), consistent with high-quality spatial chromatin accessibility profiles. Peak calling identified a total of 195,115 peaks in Black (mean 39,023 peaks/sample) and 206,996 in White men (mean 41,399 peaks/sample) (**Fig. S2B**). In both groups, accessible regions were enriched near TSS (**Fig. 1E**) and fragment size distributions displayed the expected nucleosomal periodicity, including nucleosome-free fragments (<100bp) and mono- and dinucleosome-protected fragments (∼200 and ∼400 bp, respectively) (**Fig. 1F)**. Overall, these quality metrics are consistent with published spatial ATAC-seq datasets and support robust chromatin accessibility profiling across samples.

### High-Resolution Mapping of Cellular Composition in Prostate Cancer via Spatial ATAC-seq

We next generated a unified peak-by-spot accessibility matrix using a common reference peak set across all tissue sections and performed dimensionality reduction using uniform manifold approximation and projection (UMAP). Unsupervised Louvain clustering analysis identified seven reproducible chromatin accessibility cluster (**Fig. 2A** and Fig. **S3A-B**). The overall cluster distribution was comparable between tumors from Black and White patients (**Fig. 2B** and **Fig. S3C**) with only modest differences in cluster abundance, including enrichment of cluster 3 in White tumors and clusters 4 and 7 in Black tumors. We further assessed cluster robustness using kMM (k-nearest-neighbor Mutual Mapping) analysis, which compares local neighborhood relationships across cluster assignments. Both origin and host views confirmed the spatial coherence and distinct composition of individual clusters, supporting robust separation of chromatin states across tissue sections (**Fig. S3C-D**). Cluster 7 was the only exception, showing partial mixing with clusters 5 and 6, consistent with its less well-defined segregation in the UMAP embedding.

**Figure 2:**
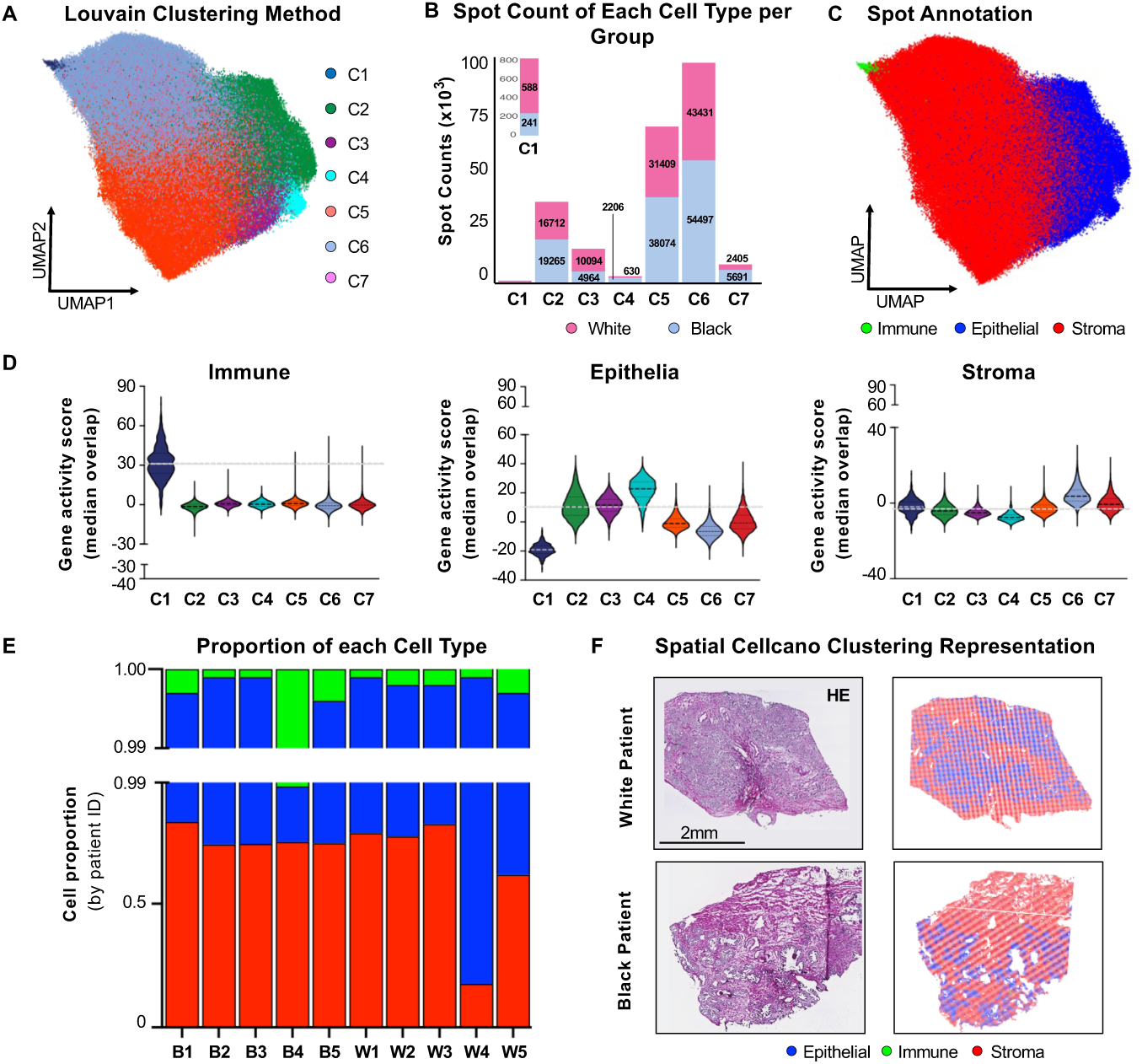
High-Resolution Mapping of Cellular Composition in Prostate Cancer via Spatial ATAC-seq. **A.** UMAP visualization of the seven Louvain clusters identified using Seurat’s workflow. **B.** Bar plot showing the number of spot assigned to each clusters across White and Black patient samples. **C.** UMAP visualization of the major cell types within the PCa tumor microenvironment (Epithelial, immune, stromal), as predicted using the Cellcano machine learning framework trained on a pan-cancer single-cell ATAC-seq atlas (Zhang K, et al. 2021). **D.** Violin plots representing gene activity scores for canonical immune, epithelial, and stroma marker genes stratified by cluster. **E.** Bar plot representing the proportion of each predicted cell type per patient sample (B, Black; W, White). **F.** Representative H&E staining (left, Scale bar= 2mm) and corresponding spatial maps of the predicted cell-type annotations from a White and Black Patient.

To annotate cell identities, we applied the machine learning classifier tool Cellcano (34) trained on a reference scATAC-seq dataset spanning 30 adult different human cancer tissue types (35) followed by independent validation using canonical marker accessibility and WebCSEA (<u>Web</u>-based <u>C</u>ell-type <u>S</u>pecific <u>E</u>nrichment <u>A</u>nalysis of Genes) enrichment analysis (36) (reference dataset Henry et al. (37)) (**Fig. 2C-E; Fig. S3F-H; Fig.S4; Tables S2-S5**). Together, these complementary approaches consistently identified one immune cluster (cluster 1), three epithelial clusters (clusters 2-4), and three stromal clusters (clusters 5-7). Cell identities were further supported by chromatin accessibility at canonical lineage markers, including KRT8, KRT18, and NKX3-1in epithelial cells, RUNX3 and PTPRC in immune cells, and TAGLN, DCN, ACTA2, MYL9, and DES in stromal cells (**Fig. S5**).

To assess global ancestry-associated differences in chromatin accessibility, we next performed pseudobulk analysis by aggregating reads across all spatial ATAC-seq profiles. Fragment counts per spatial barcode were comparable between groups, indicating similar sequencing depth. Although tumors from Black patients exhibited a modest increase in promoter-accessible regions (584 versus 512 peaks), the genomic distribution of accessible regions was highly similar between groups, with most peaks mapping to intronic and distal intergenic regions (**Fig. S6**). Likewise, the relative abundance of epithelial, immune, and stromal populations was comparable between Black and White tumors (**Fig. 2E**; **Fig. S3G**). These findings indicate that ancestry-associated differences are unlikely to reflect global changes in cellular composition or chromatin accessibility, supporting subsequent analyses of cell type-specific regulatory programs.

### Spatial inference of copy-number alterations reveals clonal architecture in prostate cancer

Tumor spots were independently identified by integration of pathologist-based histological annotation, tumor-associated gene activity scores, and copy-number variation (CNV) profiles, thereby providing complementary morphological, transcriptional, and genomic criteria for tumor assignment. CNVs from spatially resolved ATAC profiles using the EpiAneufinder (EpiAn) algorithm were inferred, extending single-cell ATAC-seq-based CNV inference to spatially resolved datasets (38). EpiAn identified CNV patterns across individual spatial barcodes and classified genomic regions into gain, loss, or neutral states. This analysis revealed eight distinct karyotype clones (A1-A8) (**Fig. 3A**) with spatial distribution overlaying histologically annotated tumor regions (**Fig. 3B**). Notably, clone 6 (A6) was detected at low frequency across all samples (**Fig. 3C**) and exhibited minimal detectable CNV alterations and the lowest tumor-associated gene activity score (**Fig. 3C-E**). A6 was therefore classified as non-malignant and all remaining clones (A1-A8) were classified as malignant. This strategy enabled us to distinguish malignant epithelial regions. Therefore, downstream analyses were restricted to the malignant spots.

**Figure 3.**
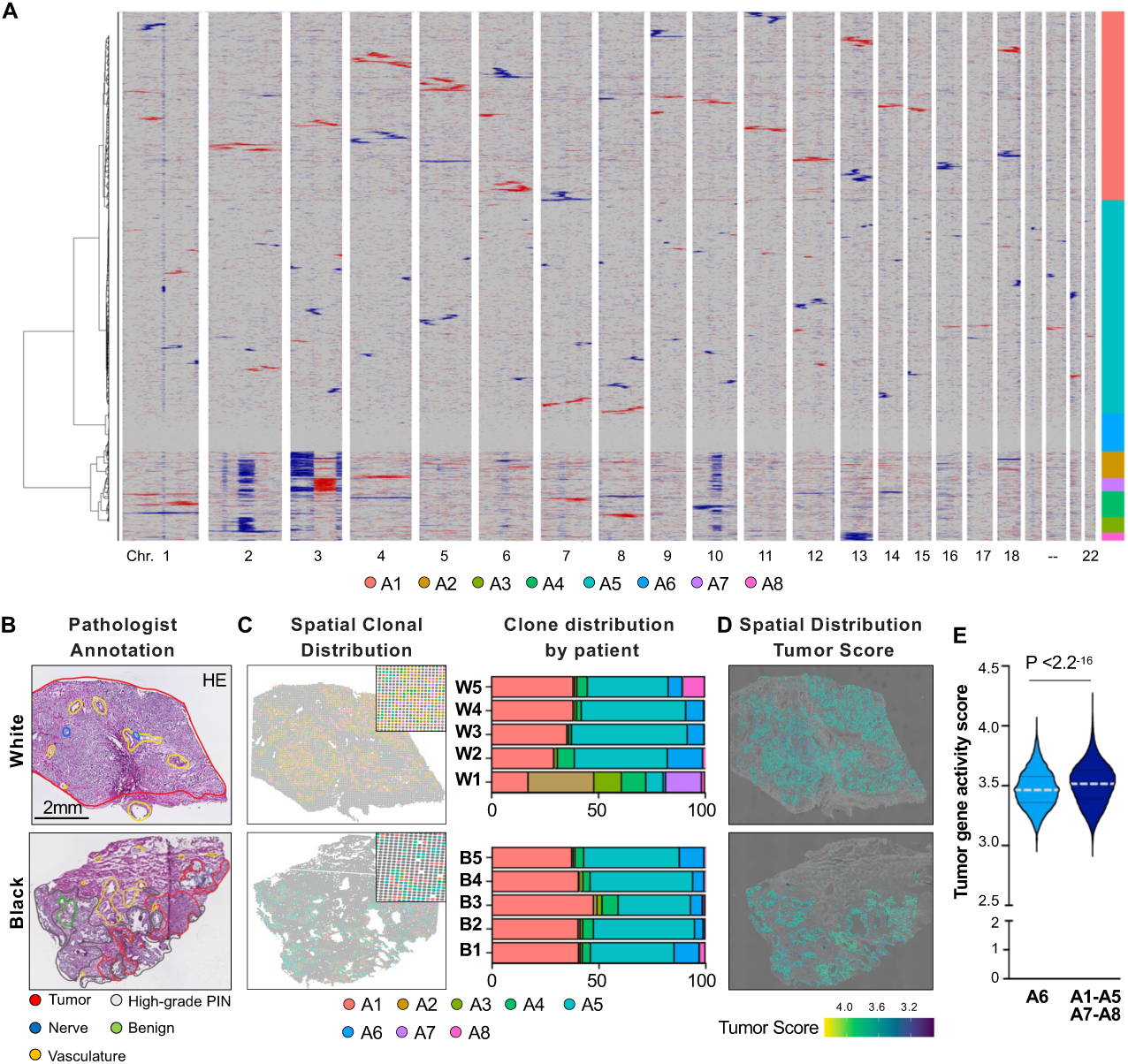
Race-associated copy number alterations identified by spatial ATAC-seq. **A.** Inferred copy number variation (CNV) profiles from spatial ATAC-seq data identifying eight distinct karyotypic clones (A1–A8). Each row represents a single cell, and each column represents a chromosome. Karyotypic clones are color-coded and indicated in the sidebar. **B.** Representative H&E-stained sections from a White and a Black patient, annotated by a pathologist. Red indicates tumor tissue, gray indicates high-grade prostatic intraepithelial neoplasia (HGPIN), green indicates benign tissue, and yellow indicates vasculature. Scale bar, 2 mm (left). **C.** Spatial distribution of karyotypic clones in representative White (W) and Black (B) patients, accompanied by bar plots showing the relative abundance of each clone (A1-A8) across patients. **D.** Representative spatial distribution of the tumor gene activity score in a White and a Black patient. **E.** Violin plot comparing the tumor gene activity score between Clone A6 (lacking detectable copy number alterations) and all remaining clones (A1-A5 and A7-A8). Statistical significance was assessed using the Wilcoxon rank-sum test.

### Spatial epigenomic profiling reveals heterogeneous epithelial states in prostate cancer

The unsupervised Louvain analysis resolved three major epithelial chromatin states (E2-E4) (**Fig. 4A**), revealing heterogeneity in the regulatory landscape of malignant epithelial regions. The distribution of these states formed a continuous landscape rather than sharply separated populations. Consistent with this organization, pseudotime analysis identified a trajectory connecting E2-E4, with coordinated changes in epithelial lineage-associated programs (**Fig. 4B**). E2 regions were predominantly positioned at earlier pseudotime values, whereas E3 and E4 occupied two distinct terminal branches. These findings identify E2-E4 as distinct chromatin states within a continuous epithelial regulatory landscape.

**Figure 4:**
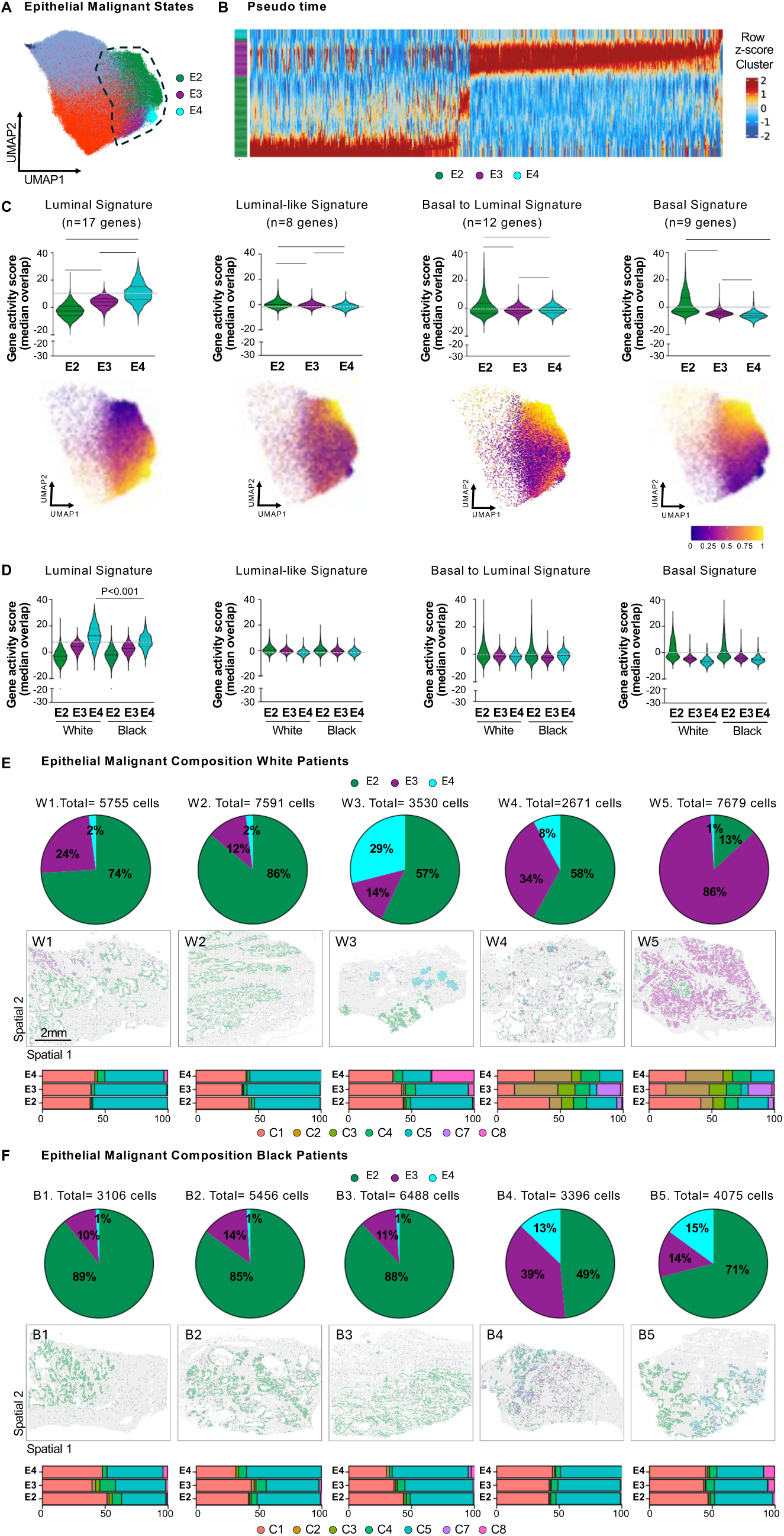
Epithelial annotation identified in Black and White Prostate Cancer. **A.** UMAP highlighting the three epithelial malignant states (E2, E3, and E4). **B.** Pseudotime analysis representing spline-fitted average Z-scores of progenitor score in the epithelial malignant states**. C.** Violin plots and UMAPs representing the gene activity score for luminal, luminal-like, basal to luminal and basal signatures in the three epithelial malignant states. **D.** Violin plots representing the gene activity score for luminal, luminal-like, basal to luminal and basal signatures segregated by race in the three epithelial malignant states. **E.** Pie chart representing proportion of each epithelial state stratified by patient in the White group (up). Representative spatial images for the epithelial state stratified by patient in the White group (middle). Bar plot representing the proportion of each tumoral clone in the different epithelial state stratified by patient in the White group (down). **F.** Pie chart representing proportion of each epithelial state stratified by patient in the Black group (up). Representative spatial images for the epithelial state stratified by patient in the Black group (middle). Bar plot representing the proportion of each clone in the different epithelial state stratified by patient in the Black group (down).

We next characterized the lineage-associated programs represented across E2-E4. Luminal, luminal-like, basal-to-luminal, and basal-associated programs showed differential representation across the three states (**Fig. 4C**). E3 and E4 were predominantly enriched for luminal programs, whereas E2 displayed a broader profile combining luminal and basal-associated features, together with enrichment of a basal-to-luminal transition signature. Thus, E2-E4 differed in the representation of epithelial lineage-associated and translational regulatory programs.

The abundance and spatial organization of E2-E4 varied across patients (**Fig. 4E-F**). E2 was the predominant state in most tumors in both cohorts, whereas E3 or E4 represented a larger fraction in individual tumors. Spatial mapping further showed patient-specific organization of these states within tumor regions (**Fig. 4E-F**). E4 regions from White tumors showed significantly higher luminal gene-score activity than E4 regions from Black tumors (**Fig.4D**). Together, these analyses demonstrate substantial inter-patient and intra-tumor heterogeneity in the chromatin landscape of PCa, characterized by coordinated shifts in epithelial lineage-associated and transitional regulatory programs.

### Population differences in epithelial regulatory programs are associated with differential CTCF-associated chromatin states

To define population-associated differences in the regulatory landscape of malignant epithelial states, we first performed motif enrichment analyses within each epithelial chromatin state. In tumors from White patients, E2-E3 states showed enrichment for luminal lineage-associated regulatory factors, including CTCF, AR, and NR3C1, with additional enrichment for SMARCC1 in E2 and NRF1 and GABPA in E3 (**Fig. 5A**), AP-1 family motifs, including JUN, JUNB, JUND, FOSB, and FOS, were also enriched across E2-E4, although the specific motifs and magnitude of enrichment varied between epithelial states. In contrast, epithelial states from Black patients showed greater enrichment for developmental and phenotypic plasticity-associated regulators, including ZEB1, TCF4, ID3/4, and MESP1/2 in E2-E3, as well as KLF family members in E4, FOSL2 in E3, and distinct FOX-family motifs in E2 (**Fig. 5A)**. These findings indicate that malignant epithelial states from the two populations retain distinct regulatory features, with Black tumors showing greater representation of developmental and plasticity-associated transcriptional programs.

**Figure 5.**
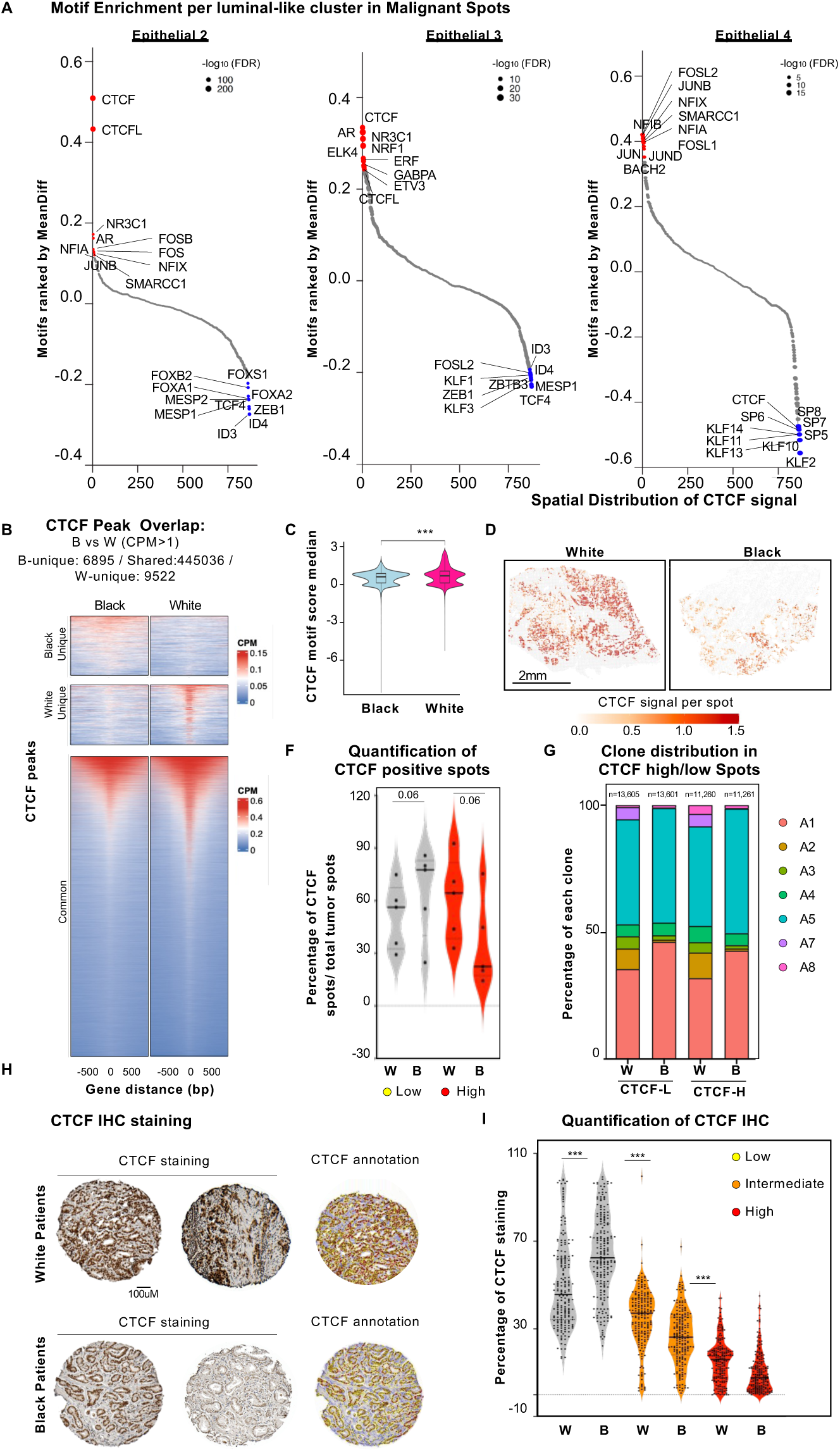
Epigenetically defined epithelial states are associated with distinct regulatory programs in Black and White prostate cancer. **A.** Differential motif enrichment elbow plot within malignant epithelial chromatin states (Epithelial 2–4), stratified by patient group. Motifs are ranked by mean difference in enrichment between groups; top 10 enriched motifs with are associated with luminal, lineage-associated, and developmental/plasticity-associated motifs are indicated. **B.** Distribution of CTCF motif accessibility scores in malignant epithelial regions from Black and White tumors. **C.** Violin plot representing the differential motif analysis of malignant epithelial regions, showing CTCF and CTCF-like motifs among the strongest population-associated features. **D.** Overlap of CTCF-associated accessible regions between White and Black tumors, showing that most regions are shared, with a small fraction showing group-specific enrichment. **E.** Representative spatial distribution image of CTCF-associated signal across malignant epithelial regions in White and Black tumors. **F.** Quantification of the proportion of malignant epithelial spots classified as CTCF-high or CTCF-low across White and Black tumors. **G.** Distribution of epithelial malignant clones within CTCF-high and CTCF-low spots across White and Black tumors. **H.** Representative CTCF immunohistochemistry and corresponding CTCF annotation in an independent prostate cancer tissue microarray from White and Black patients. **I.** Quantification of CTCF protein staining, showing the distribution of CTCF-low, intermediate, and high tumors across patient groups. Statistical significance is indicated as shown.

Among the differentially enriched motifs, CTCF was the most consistent population-associated differences across the epithelial states (**Fig. 5A**). Importantly, this difference was not driven by extensive remodeling of the underlying CTCF-associated regulatory landscape. Most CTCF-accessible regions were shared between groups, with only a small fraction showing group-enriched accessibility (**Fig. 5B**). We therefore examined CTCF-associated chromatin accessibility across malignant epithelial spots. CTCF motif accessibility was significantly higher in malignant epithelial regions from White tumors than in those from Black tumors (**Fig. 5C-G**), and the clone distribution was not dependent on CTCF-high (CTCF-H) or CTCF-low (CTCF-L).

To independently validate this observation at the protein level, we assessed CTCF expression by immunohistochemistry in an independent tissue microarray comprising of 99 localized PCa specimens (50 White and 49 Black patients). Consistent with the spatial ATAC-seq analysis, tumors from Black patients exhibited a significantly higher proportion of CTCF-low (L-CTCF) cases than tumors from White patients (**Fig. 5H-I**), providing evidence that reduced CTCF-associated regulatory activity is a feature of malignant epithelial states in Black tumors.

## Discussion

Our study provides a spatial epigenomic characterization of primary PCa using radical prostatectomy specimens from Black and White patients treated within a healthcare system with equal access to care. By applying spatial ATAC-seq, we mapped chromatin accessibility while preserving tissue architecture, enabling the identification of distinct cellular compartments and malignant epithelial states within individual tumors. Spot-type annotation and CNV profiles identified malignant epithelial regions, providing a framework to compare chromatin states within the tumor compartment. This spatial approach is particularly relevant to PCa, where substantial regulatory heterogeneity can be obscured by bulk profiling (3–7) (39).

Within the specimens the overall immune, stroma and epithelial density did not differ significantly between Black and White patients, consistent with previous studies (40–43). Yet in the malignant epithelial compartment, we identified distinct chromatin accessibility landscapes between the two groups. This suggests that the observed differences are not simply associated to differences in the representation of malignant epithelial regions but instead reflect variation in their regulatory states. Importantly, spatial profiling allowed these features to be resolved within individual tumor regions rather than averaged across heterogeneous specimens. Although the current cohort comprises 10 patients, these findings establish a framework for examining population-associated chromatin states within defined malignant epithelial compartments in larger cohorts.

Focusing on malignant epithelial cells, a reduced accessibility of CTCF motif in tumors from Black patients was particularly prominent and was accompanied by reduced CTCF protein expression in an independent tissue microarray. CTCF is a major regulator of tree-dimensional genome organization (44), and both CTCF expression and motif accessibility have been linked to EMT (45, 46). Our spatial ATAC-seq analysis and immunohistochemistry provide complementary views of this regulatory axis: motif accessibility reflects the chromatin environment surrounding potential CTCF binding sites, while reduced protein expression provides evidence of altered CTCF abundance. Together, these findings identify CTCF associated regulation as a prominent feature distinguishing the chromatin landscapes of tumors from Black and White patients. Whether these differences are accompanied by altered CTCF occupancy or higher-order chromatin organization remains an important question for future studies.

Beyond CTCF associated features, tumors from Black patients showed greater enrichment of motifs linked to EMT and inflammatory programs, placing the CTCF finding within a broader chromatin landscape associated with epithelial plasticity and inflammatory signaling. The coexistence of reduced CTCF associated accessibility with enrichment of these regulatory programs suggests that differences in chromatin regulation may contribute to distinct epithelial malignant states. This observation is particularly relevant in the context of phenotypic plasticity in localized PCa, before the emergence of advanced or treatment-refractory disease. Although phenotypic plasticity has been extensively studied in metastatic and therapy-resistant PCa (18, 47), evidence from both experimental mouse models and human tumors indicates that phenotypic plasticity can emerge earlier in disease. In primary PCa, combined *Pten:Trp53* loss induces heterogeneous epithelial states and EMT promoting invasive behavior (16, 48–50). Consistent with this concept in human disease, studies of localized PCa have shown that pre-existing genomic alterations, including *TP53* and *PTEN* alterations, are associated with differential response to androgen-pathway inhibition (51, 52). Our findings extend this concept to human localized PCa by identifying spatially resolved chromatin features associated with epithelial plasticity in the context of race. While our data do not establish a direct mechanistic relationship between CTCF and the EMT or inflammatory programs, the convergence of these chromatin features highlights CTCF associated regulation as a candidate component of the regulatory landscape underlying epithelial plasticity. Integrating chromatin accessibility with CTCF occupancy, transcriptional states, and functional measurements will be important to define how these regulatory features relate to epithelial state and tumor behavior.

Overall, our findings show that population-associated heterogeneity in localized PCa extends to chromatin regulation within the malignant epithelial compartment. Reduced CTCF associated accessibility and protein expression, together with enrichment of inflammatory and EMT-associated regulatory elements, identify distinct features of the chromatin landscape between Black and White patients. Rather than defining a population-specific tumor phenotype, these findings indicate that chromatin regulatory states represent an additional dimension of PCa heterogeneity and identify epithelial plasticity as a potential feature of this regulatory landscape. Larger, integrated studies will be important to establish the reproducibility of these states and determine their relationship to tumor evolution and clinical behavior.

## Material and Methods

### Spatial ATAC-seq on frozen prostate cancer

Frozen prostate cancer samples were embedded in OCT within cryomolds by adding small volumes of OCT over dry ice to minimize tissue thawing. Ten prostate cancer samples were analyzed (N = 5 White and N = 5 Black). Clinicopathological characteristics are provided in **Fig. 1B**. Tissue sections (7 μm) were prepared using a cryostat and collected onto SuperFrost Plus charged slides (Fisher Scientific, Cat. No. 12-550-15). Sections were briefly thawed to ensure adherence to the slides and immediately returned to the −20°C cryostat chamber until sectioning was completed. Slides were subsequently transferred to AtlasXomics for spatial ATAC-seq processing.

Spatial ATAC-seq was performed using the AtlasXomics AXO-0303 workflow. Briefly, tissue sections were fixed for 5 min in 0.2% paraformaldehyde, treated with glycine, and allowed to dry. Tissue permeabilization was performed with 0.1% NP-40 for 15 min, followed by Tn5 transposition for 30 min. Spatial barcoding adapters containing ligation linkers were introduced through microchannels and ligated to the Tn5-generated DNA ends through successive rounds of ligation. Sections were imaged to enable spatial registration of barcoded accessible chromatin with tissue morphology. A barcoded tissue mosaic was generated, followed by reverse cross-linking to release DNA fragments. DNA fragments were subsequently PCR-amplified and processed for sequencing library preparation. Libraries were sequenced using 150-bp paired-end sequencing on a NextSeq 2000 with 15% PhiX. Sequencing depth was up to 300 million reads per sample.

### Processing of spatial ATAC-seq data

Spatial ATAC-seq data were processed by AtlasXomics as previously described (53). Sequencing reads were aligned to the human GENCODE reference genome (v38) using Chromap, and PCR duplicates were removed [91]. Fragment files containing the genomic coordinates of sequenced DNA molecules associated with spatial barcodes were generated for downstream analysis.

Downstream analyses were performed using an ArchR-based workflow (ArchR v1.0.2; R v4.4.1) (54). Raw fragment files generated by the AtlasXomics platform were converted into Arrow files using the createArrowFiles() function with a 5-kb tile size. Quality control was performed at the spatial-pixel level. Pixels were retained based on a minimum TSS enrichment score of 2 and the predefined upper limit for fragment counts. Spatial metadata, including Cartesian coordinates and tissue-mask designations (on-tissue versus off-tissue), were extracted from the AtlasXomics tissue positions_list.csv file and incorporated into the ArchR environment. Only pixels classified as on-tissue by paired imaging analysis were retained for downstream analyses.

### Dimensionality reduction, batch correction, and clustering

Latent semantic indexing (LSI) was performed on the 5-kb TileMatrix using addIterativeLSI() in ArchR. Batch correction across samples was performed using Harmony integration through addHarmony() applied to the LSI low-dimensional representation. Two-dimensional embeddings were generated using uniform manifold approximation and projection (UMAP) with addUMAP(). Unsupervised clustering was performed using Louvain community detection with addClusters() on the Harmony-corrected dimensions at a resolution of 0.3.

### Cell-type annotation

Cell-type identities were assigned using correlation-based prediction implemented in the deCS R package (v1.0.0) (55). Two reference atlases were used: (i) the Tabula Sapi-ens multi-tissue atlas (56) and (ii) a published prostate single-cell RNA-seq reference dataset (37). For each spatial spot (or cluster), average gene activity profiles were correlated (Spearman) with reference cell-type expression signatures, and the top-correlated cell type was assigned as the predicted label.

### CNV inference and tumor–normal classification

Copy-number profiles were inferred using epiAneufinder (38) on the hg38 reference genome with a 700-kb genomic window size. CNV states were represented as loss, neutral, or gain, resulting in eight CNV-defined clones. To summarize large-scale aneuploidy, two metrics were calculated for each spatial pixel: (i) a global CNV burden score, defined as the mean absolute deviation from the neutral state across genomic bins, and (ii) a chromosome-level alteration score, defined as the mean fraction of altered bins across the three most altered chromosomes.

Tumor-like and normal-like CNV profiles were initially identified in an unsupervised manner using a two-component Gaussian mixture model fitted to z-scored CNV features. The component with the higher mean CNV burden was designated as tumor-like, and the posterior probability of membership in this component was used as a continuous tumor probability score. To reduce spatial noise, tumor probabilities were smoothed within each sample using k-nearest neighbors based on tissue coordinates (k = 8). The final tumor probability was calculated as the mean of the raw and neighbor-averaged probabilities. For calibration and validation, a set of 354 manually reviewed cells (257 tumor-like, 97 normal-like) with pathologist-confirmed phenotype was used to select a decision threshold by ROC analysis (Youden criterion; threshold = 0.241). Cells were classified as tumor-like if their final probability exceeded the threshold and as normal-like otherwise.

### Peak calling and accessibility quantification

Accessible chromatin peaks were identified using MACS2 v2.2.9 through ArchR. Group-wise pseudobulk replicates were generated using addGroupCoverages(), and reproducible peaks were identified across replicates using addReproduciblePeakSet() to generate a consensus peak set. A sparse peak-by-pixel count matrix was subsequently generated using addPeakMatrix() for downstream analyses.

### Differential accessibility analyses

Differentially accessible peaks were identified using getMarkerFeatures() in ArchR applied to the PeakMatrix. Statistical testing was performed using the Wilcoxon rank-sum test with bias correction for TSS enrichment and sequencing depth, represented as log10-transformed fragment counts. Multiple testing was controlled using the Benjamini–Hochberg procedure. Peaks were considered differentially accessible at FDR < 0.05 and |log2FC| ≥ 0.1.

### Sample-level pseudobulk quantification

Statistical comparisons between racial groups were performed at the sample level, with individual patients/samples treated as biological replicates. For race-level comparisons, fragment counts were aggregated across spatial pixels within each sample and peak to generate a sample-by-peak pseudobulk count matrix. Counts were normalized within each sample to counts per million (CPM):

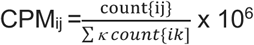

Peaks were retained if they had at least five raw counts in at least two samples. A peak was considered supported within an individual sample when CPM ≥ 1. Peaks were subsequently categorized as group-unique, when supported exclusively among samples from one racial group, or shared, when supported in at least one sample from each group. Overlap patterns were visualized using Venn diagrams.

### Motif annotation and chromVAR deviations

Transcription factor (TF) motifs were annotated within the consensus peak set using CIS-BP through addMotifAnnotations(). Background peaks matched for GC content and accessibility were generated using addBgdPeaks (). Per-spot motif accessibility wasquantified using chromVAR deviation scores computed with addDeviationsMatrix ().

### Differential motif activity

Differential motif activity was assessed using getMarkerFeatures() applied to the motif deviation matrix (MotifMatrix), using Wilcoxon rank-sum with correction for technical covariances including TSS enrichment and sequencing depth (log10-transformed fragment counts) through bias-matched background selection.

Motifs were considered significant at FDR < 0.05 with mean deviation difference ≥ 0.1. Unless otherwise stated, statistical testing was performed on non-imputed values; imputed values were used for visualization.

### Gene activity and pathway analysis

Gene activity scores were calculated from chromatin accessibility using addGeneScoreMatrix() in ArchR, which aggregates accessibility signal across gene bodies and promoter-proximal regions using distance-based weighting relative to transcription start sites. Imputation weights were calculated using addImputeWeights() based on the Harmony-corrected low-dimensional embedding, and an imputed gene activity matrix was generated using imputeMatrix() applied to the GeneScoreMatrix.

Unless otherwise indicated, statistical analyses were performed using non-imputed gene activity values, whereas imputed values were used for visualization and pathway-level scoring. Imputed gene activity matrices were imported into Seurat (v5.1.0) as an assay for downstream visualization. Pathway signature scores were calculated for each spatial pixel by aggregating imputed gene activity values across genes included in the corresponding pathway gene sets (gene sets specified in Data Availability).

### Computational environment

All computational analyses were performed on the NIH Biowulf high-performance computing cluster using R v4.4.1. Core packages included ArchR v1.0.2, Seurat v5.1.0, deCS v1.0.0, ggplot2 v3.5.1, and ComplexHeatmap v2.20.0. Detailed software versions, analysis parameters, and scripts are available in the accompanying repository.

## Acknowledgments

This study was supported from a Department of Defense Idea Development Award (HT94252510166 to LE and BG).

## Author Contributions

Conceptualization: LE, BG, NC

Methodology: LE, BG, KL-H, SS, RS, GJ, NC, ES

Investigation: LE, BG, KL-H, SS, RS, GJ, AA, AW, ES

Analysis: LE, BG, KL-H, SS, RS, GJ, NC, AA, AW, ES

Visualization: LE, BG, KL_H, SS, RS, NC

Funding acquisition: LE, BG

Project administration: LE, BG Supervision: LE, BG

Writing – original draft: LE, BG, KL-H, SS, RS, CG

